# Infant formula composition is linked to distinct patterns of early gut colonization

**DOI:** 10.64898/2026.11.20.768025

**Authors:** Ioanna Fountouli, Yuvashankar Jayaprakash, Twan Keizer, Trishla Sinha, James A. D. Docherty, Lifelines Next Cohort Study, Alexander Kurilshikov, Hermie Harmsen, Alexandra Zhernakova, Cyrus A. Mallon

## Abstract

Early-life nutrition shapes the gut microbiota and can have long-term health implications. While infant formula feeding is widespread, it remains unclear how variations in its nutritional components shape microbiome colonization. Here, we reconstructed 62 components across the 17 infant formulas consumed during the first three months of life in 240 exclusively formula-fed infants in the Lifelines-NEXT birth cohort. Formula composition varied widely and several micronutrients, including vitamin E and iodine, were associated with species richness. Components were also associated with species-level colonization patterns. Higher iron content was associated with reduced *Bifidobacterium breve* and *Bifidobacterium bifidum* and increased *Clostridium paraputrificum*. Follow-up *in vitro* growth assays on infant-derived stool isolates showed that *Bifidobacterium breve* and *Bifidobacterium bifidum* tolerated iron limitation, while *C. paraputrificum* did not, consistent with the association signals. These findings show that infant formulas are not all alike and differences in component concentration affect microbiome colonization in early-life.

**Highlights:**

- Infant formula composition varies widely and shapes early-life gut microbiota
- Formula components modulate species richness and select for distinct bacterial taxa
- Higher iron is associated with reduced Bifidobacteria and increased Clostridia
- Bifidobacteria isolated from infant stools are adapted to low iron availability

## Introduction

Early-life nutrition is a major determinant of infant development and gut microbiome colonization. Although breast milk is considered the optimal source of infant nutrition, infant formula is widely used when breastfeeding is not possible or sufficient ^1,2^. Indeed, it is estimated that only 44% of infants 0–6 months are exclusively breastfed worldwide ^3^. While considerable research has focused on how breast milk components influence microbial colonization and infant development, less is known about the impact of specific macro- and micronutrients in formula. Formula composition differs from that of breast milk and varies across products in both nutrient concentrations and the inclusion of bioactive ingredients. These differences may contribute to variation in microbial colonization during a period when the gut microbiome and host immune and metabolic systems are developing ^1,4–6^. As such, understanding how formula composition relates to the infant gut microbiome is important for informing efforts to improve infant nutrition.

To mimic the effects of breast milk on the developing gut microbiome, infant formulas are increasingly supplemented with bioactive ingredients. Breast milk bioactives support infant development by promoting colonization by commensal bacteria, inhibiting pathogens, and supporting mucosal immune development ^7,8^. Formula supplementation aims to reproduce some of these functions and their potential benefits for developmental outcomes. Common additions include prebiotics such as galacto-oligosaccharides (GOS) and fructo-oligosaccharides (FOS), probiotic strains of *Bifidobacterium animalis* subsp. *lactis* and *Lactobacillus rhamnosus*, and human milk oligosaccharides (HMOs) such as 2′-fucosyllactose ^9,10^. Much of the research on formula composition and the infant microbiome has narrowly focused on these selected ingredients, while the role of broader variation in macro- and micronutrient concentrations is less well characterized.

Despite these developments, differences in gut microbiome composition persist between formula-fed and breastfed infants. Our previous work showed that formula-fed infants exhibit greater bacterial species diversity^11^ and lower abundances of infant-associated *Bifidobacterium* species^12^. Formula feeding has also been associated with higher relative abundances of *Clostridium* and taxa within the Enterobacteriaceae family ^13–15^. Feeding-associated differences in the infant microbiome have also been linked to variation in growth and metabolic outcomes ^6,16^. However, the mechanisms driving these microbiome differences remain poorly understood. In particular, it is unclear how variations in specific components of infant formula influence microbial colonization and competitive interactions in early-life. We hypothesized that differences in the concentrations of specific formula components, including macronutrients, vitamins, and minerals, would affect competitive dynamics among bacterial species, thereby shaping gut microbiota composition. Understanding how component variability in formula relates to microbiome characteristics is essential for optimizing formula composition and promoting healthy microbiome development in formula-fed infants.

We investigated this hypothesis using data from the extensively characterized Lifelines NEXT (LLNEXT) cohort. We conducted a longitudinal analysis of 470 faecal samples from 240 exclusively formula-fed infants collected at four time points during the first three months of life. We employed mixed regression models to assess the associations of 51 formula components with microbial diversity and specific bacterial taxa. Finally, we validated key findings using *in vitro* growth experiments examining how iron availability affects bacterial growth.

## Methods

### EXPERIMENTAL MODEL AND STUDY PARTICIPANT DETAILS

#### Lifelines NEXT cohort study

##### Study design

The samples for this study were obtained from the LLNEXT cohort, a mother–infant cohort designed to study the effects of intrinsic and extrinsic predictors on health and disease during early life^17^. From 2016 to 2023, 1,452 pregnant women were recruited for LLNEXT. Mothers were monitored pre- and post-pregnancy, and infants were monitored during their first year of life. This included collection of biomaterial and data on medical, social, and environmental factors collected via questionnaires. The present study includes the 240 LLNEXT infants who were exclusively formula-fed, monitored at week 2 (W2), month 1 (M1), month 2 (M2), and month 3 (M3) of life. Later timepoints were excluded to eliminate bias in analyses due to solid-food introduction. Basic cohort summary statistics are shown in Figure 1A.

##### Ethical approval

The Lifelines NEXT cohort study has been approved by the Ethics Committee of the University Medical Center Groningen (METC UMCG METc2015/600). Written informed consent was obtained from participants or their parents/legal guardians before participation.

##### Bacterial Strains

Bacterial strains used in this study were isolated from faecal samples taken between W2 and M6 from infants who were part of the LLNEXT cohort ^11^. Isolates included *Bifidobacterium breve*, *Bifidobacterium bifidum* and *Clostridium paraputrificum*. Bacteria were cultured in the following media: Yeast Casitone Fatty Acids Broth (YCFA) supplemented with 20 mM Lactose (YCFA-L) and 0.05% (w/v) L-cystine. All bacteria were grown under anaerobic conditions (90% nitrogen, 5% hydrogen, 5 % carbon dioxide) at 37°C.

### METHOD DETAILS

#### Samples

##### Infant faecal samples

For this study, 470 infant faecal samples were collected from 240 infants at W2 (*n* = 64), M1 (*n* = 124), M2 (*n* = 130), and M3 (*n* = 152). Infant faecal samples were collected by parents/guardians and processed, as previously described ^11^. Briefly, samples were frozen at home, transported to the University Medical Center Groningen in portable freezers, and stored at -20°C (short-term) or -80°C (long-term) until analysis.

##### Phenotypic data

Infant feeding mode and formula type were obtained from food frequency questionnaires filled in by parents at W2, M1, M2, and M3. Infants were included if they were exclusively formula-fed and received a single formula type at each respective timepoint. In total, 17 formula brands were consumed by the participants in our cohort. The distribution of samples across the 17 formula brands is visualized in Supplementary Figure 1, which reports both overall sample frequency per formula and sample frequency at each timepoint. Formula milk composition data were collected from publicly available nutritional tables on manufacturers’ websites (accessed March 2025). A total of 62 components present in formula milk were recorded, along with their concentrations per 100ml of prepared formula. In this study, formula components refer to all labelled constituents of the infant formulas, including macro- and micronutrients and added bioactive ingredients (e.g. FOS, GOS). Delivery mode was recorded from birth cards completed by midwives or obstetricians at the hospital.

##### Faecal DNA isolation and shotgun metagenomic sequencing

Microbial DNA was isolated from 0.2–0.5 g faecal material using the QIAamp Fast DNA Stool Mini Kit (Qiagen, Germany) and the QIAcube (Qiagen). Shotgun metagenomics sequencing was performed by Novogene (Cambridge, UK) using HiSeq 2000 or NovaSeq 6000 sequencing with 2 × 150 bp paired-end chemistry (Illumina).

##### Metagenomic Processing and Taxonomic Profiling

Bioinformatic analysis was performed using an in-house bioinformatic pipeline (https://github.com/GRONINGEN-MICROBIOME-CENTRE/gmc-mgs-pipeline). In short, adapters were first trimmed from gut microbiome sequencing reads with BBDuk (v39.01) (http://bbtools.jgi.doe.gov) and then quality trimmed using KneadData tools (v0.10.0) ^18^. The KneadData-integrated Bowtie2 tool (v2.4.2) ^19^ was used to remove reads that aligned to the human genome (GRCh38/hg38), and the quality of the processed data was examined using the FastQC toolkit (v0.11.9). Taxonomic composition of metagenomes was profiled using the MetaPhlAn4 tool with the MetaPhlAn database of marker genes mpa_vJan21 and the ChocoPhlAn Species level Genomic Bin database (202103).

##### Assessment of iron-related genes in infant-associated metagenome-assembled genomes

To investigate the presence of iron-related genes, we analyzed high-quality metagenome-assembled genomes (MAGs) constructed from infant faecal samples collected between W2 and M3. To minimise redundancy, genomes were dereplicated at strain level using dRep (v3.4.2)^20^ with an average nucleotide identity (ANI) threshold of 99.9%. All MAGs were generated from infant samples within the LLNEXT cohort^17^. For full information regarding sample processing and MAG assembly, please refer to the methods in Sinha et al. (2026) ^11^. Only MAGs with genome completeness > 90% and contamination < 5% were included, as determined by CheckM (v1.0.12, default within metaWRAP v1.3.2) ^21,22^. Coding sequence (CDS) prediction and .faa file generation was performed using Prokka (v1.14.5) ^23^. Due to the stringent quality cutoffs applied to genome assemblies, only 22 *B. bifidum* MAGs and 17 *B. breve* MAGs from formula-fed infants met the inclusion criteria, and no *C. paraputrificum* MAGs passed these thresholds. Therefore, to maximise the number of genomes available for comparative analyses, all high-quality MAGs recovered at W2–M3 were included, regardless of feeding mode. The final dataset of MAGs comprised *B. bifidum* (n = 299), *B. breve* (n = 215), and *C. paraputrificum* (n = 10) originating from at least 338 infants, although four MAGs could not be linked to a specific infant.

Iron-related genes were initially identified using FeGenie (v1.2) under default parameters ^24^. Next, we downloaded each protein sequence that comprises the experimentally characterised iron-uptake *bfeUOB* operon from *B. breve* UCC2003 ^25^ from NCBI ^26^. These proteins are the OFeT family high-affinity iron permease BfeU (WP_015438311.1), the iron-binding transporter-associated protein BfeO (WP_003831328.1), and the membrane-spanning protein BfeB (ABE94932.1). BLASTP homology searches were performed using BLAST+ (v2.13.0)^27^ against the predicted proteome of each MAG (e-value threshold of 1e^-5^). Hits that shared an amino acid identity ≥70% with query sequences were retained as true homologues. All retained hits exhibited ≥90% query coverage.

##### Bacterial strain isolation and identification

Bacterial strains were isolated in April 2025 in Groningen, Netherlands from infant stool samples (W2 to M6). Faecal samples were spread-plated onto both non-selective and selective media. *C.paraputrificum* was isolated using non-selective cultivation on YCFA agar supplemented with 20 mM lactose and 0.05% (w/v) L-cystine. *B. breve* and *B. bifidum* were isolated using Bifidobacterium selective medium consisting of Wilkins-Chalgren agar supplemented with 10 g/L 2-Fucosyl Lactose, 0.05% (w/v) L-cystine, and 100mg/L antibiotics (mupirocin or metronidazole). These plates were incubated in an anaerobic chamber (90% nitrogen, 5% hydrogen, 5 % carbon dioxide) for 72 hours. Each quadrant of the spread plate was sub-cultured onto fresh nutrient agar, and individual colonies were subsequently picked and purified. Purified colonies were identified up to species level using direct transfer method matrix-assisted laser desorption/ionization coupled to a time-of-flight mass spectrometer (MALDI-TOF smartfleX, Bruker Daltonics, Germany). Glycerol stocks of the identified isolates were prepared and stored at -80°C.

##### Iron Limitation Growth Assays

Bacteria were cultured in YCFA media supplemented with 20 mM Lactose (YCFA-L) and 0.05% (w/v) L-cystine. All bacteria were grown under anaerobic conditions (90% nitrogen, 5% hydrogen, 5 % carbon dioxide) at 37°C for 24 hours. To minimize the influence of pre-culture conditions, a starvation step was performed by washing the cells twice with PBS and incubating them in PBS for 2 hours prior to inoculation into the iron chelator YCFA-L medium. Cultures were then seeded in 96-well plates and cells were provided with the iron chelator bathophenanthrolinedisulfonic acid disodium salt hydrate (BPDS, B1375, Sigma, United Kingdom) at final concentrations ranging from 0–70μΜ. Samples were analysed using an Agilent Biotek Epoch 2 Microplate Reader (BioTek instruments, Winooski, VT, USA ), which monitors the growth of microorganisms by measuring the optical density at 600 nm (OD600) of the liquid growth medium. The experiments were run for 24 hours, with OD600 measured every 30 min, at 37°C under anaerobic conditions.

### QUANTIFICATION AND STATISTICAL ANALYSIS

#### Statistical analyses

All statistical analyses were performed and all plots created in R (version 4.4.1). For all analyses, we defined a nominal significance at p < 0.05 and statistical significance at false discovery rate (FDR) < 0.05. FDR was controlled using the Benjamini–Hochberg procedure.

#### Preparation of data for use in association studies

##### Microbiome Data Processing and Transformation

For the downstream analysis of bacterial taxa at species level, we set a relative abundance cut-off of 0.001% and a prevalence cut-off of 10%, resulting in 70 species for further analysis. Species were centre log ratio (CLR)-transformed using the geometric mean of the relative abundance of microbial taxa as the CLR denominator. Since CLR-transformation cannot be applied to zero values, we added a pseudo value of 1x10^-6^ to species abundances in these cases. To examine the presence or absence effects of microbial taxa, species were binary-transformed based on their presence or absence. In this binary transformation, abundances greater than zero were considered present (value of one) and less than zero were considered absent (value of zero). For the associations with alpha diversity, outliers were defined as values deviating ±3 standard deviations from the mean and were excluded from downstream modelling. This filtering was also applied to alpha diversity metrics (Shannon diversity and species richness).

##### Formula Data Transformation and Processing

For each formula component, we calculated the median concentration across all 17 formula brands and classified individual values as either “at/below” or “above” this median value. This binary classification enabled comparison across component concentration scales. Components not present in a given brand were treated as missing rather than assigned a value of zero and were therefore not included in the median calculation. This approach allowed us to focus on the variability in concentration across present components, rather than their absence, which could otherwise inflate analytical interpretation.

Components were included in the association analyses if they met two criteria: presence in more than 4 of the 17 formula brands, to ensure sufficient variability for modelling, and at least 10 samples per median-based group (≤ median vs. > median) at one or more timepoints. Based on these criteria, 51 of 62 components were included in the analyses. Polysaccharides, molybdenum, chromium, maltose, beta palmitate, starch, 3’-galactosyllactose, and the bacterial strains *Limosilactobacillus fermentum* and *Lactobacillus rhamnosus* GG were excluded for not meeting the first criterion. Casein was excluded for not meeting both criteria, which resulted in a substantial imbalance between the low- and high-concentration groups.

##### Associations of formula components with alpha diversity metrics and bacterial species abundance

To test the effect of formula components on Shannon diversity and species richness during the first 3 months of life, we tested this as a fixed effect in a mixed linear regression model using the package lmerTest (v3.1-3) ^28^, adding read depth, sample DNA concentration, batch effect, and timepoint as covariates and sample ID as a random effect. Given the strong effect of delivery mode on bacterial composition found in a previous study with the LL-NEXT cohort^11^, we also corrected for the effect of delivery mode. Alpha diversity was assessed using the Shannon diversity index in R package vegan (v2.6-2) ^29^. In addition to Shannon diversity, species richness was computed by summing the number of species with non-zero abundance in each sample. Tests were performed with the binary classified formula components.

*Shannon diversity index ∼ read depth + dna conc + timepoint + batch number + delivery mode +* ***formula component*** *+ (1|NEXT_ID)*

*Species richness ∼ read depth + dna conc + timepoint + batch number + delivery mode +* ***formula component*** *+ (1|NEXT_ID)*

For associations with specific bacterial species, we associated the binary-classified components with the CLR-transformed species abundance as the outcome, following the same procedure.

CLR_transformed_species_abundance ∼ read depth + dna conc + timepoint + batch number + delivery mode + **formula component** + (1|NEXT_ID)

##### Associations of formula components with bacterial species presence

To further examine the effect of formula components on the presence or absence of bacterial species during the first 3 months of life, we tested this as a fixed effect in a mixed logistic regression model using the package lme4 (v1.1-36) ^30^, adding read depth and timepoint as covariates and sample ID as a random effect. Again, given the strong effect of delivery mode on bacterial composition, we also corrected for the effect of delivery mode. We performed a single set of tests in which we associated the binary classified components with the binary transformed species presence or absence.

Binary_transformed_species_presence ∼ read depth + timepoint + delivery mode + **formula component** + (1|NEXT_ID)

## Results

### Study design and compositional variability across infant formulas

This study included data from 240 exclusively formula-fed infants, encompassing 470 faecal samples collected between 0.5 and 3 months of age and corresponding formula questionnaire data (Fig. 1A). Faecal metagenomic profiles were generated as previously described^11^ and integrated with formula composition data. We recorded a total of 62 components present in the 17 infant formula brands consumed by the participants (Fig. S1). Formula components were filtered according to the criteria described in the Methods, with 51 components retained for downstream analyses. Each component was classified as at/below or above the median concentration across the 17 formula brands (Fig. 1B). We then characterized the variability in the concentration of the components (Fig. 1C). Although total energy, fat and carbohydrate content were relatively consistent across formulas, other macro- and micronutrients showed substantial variability. Overall, 33 of 62 formula components exhibited a coefficient of variation (CV) > 25%, and 7 of 62 components showed a CV > 50% (Fig. 1C).

**Fig. 1.**
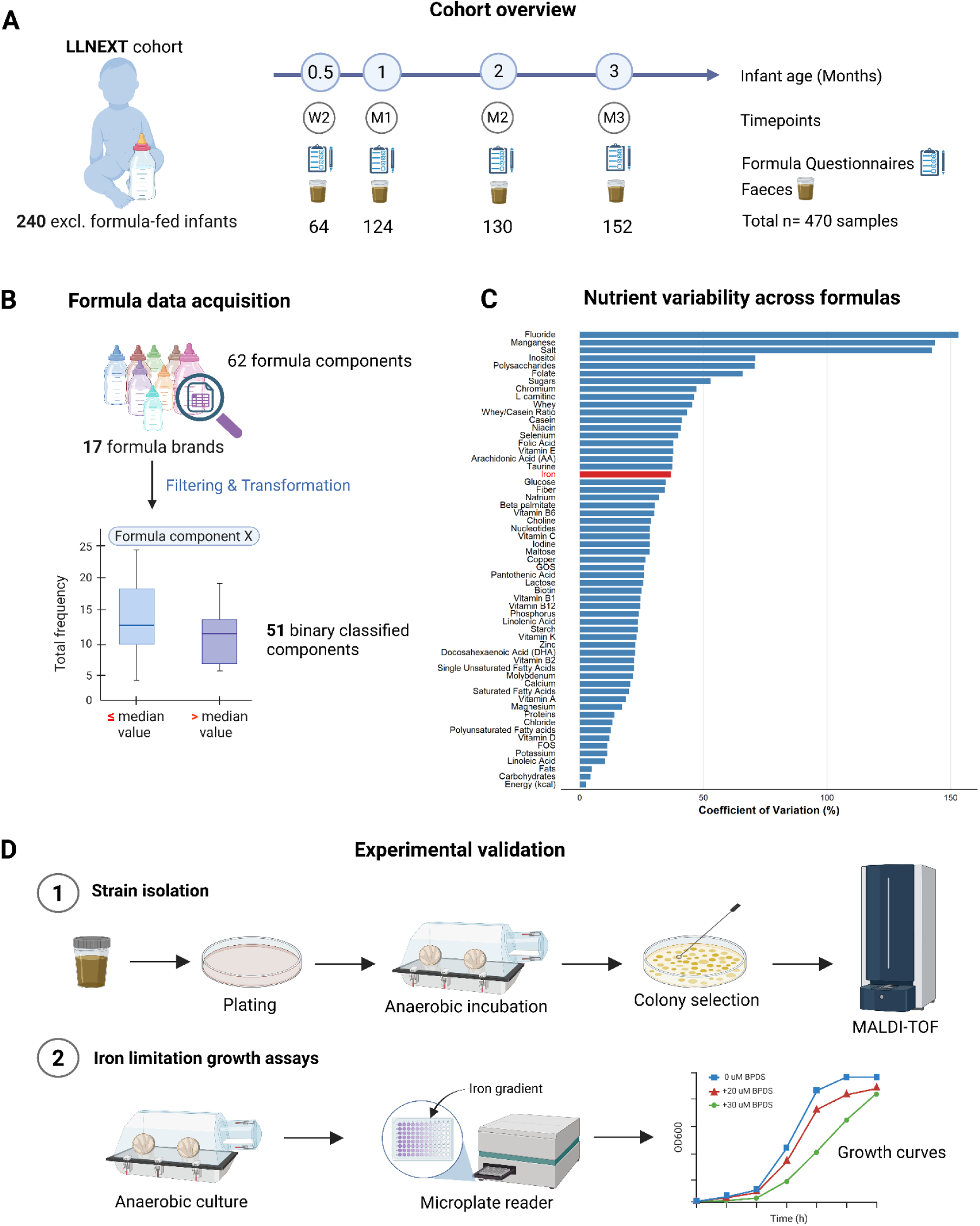
Study design and compositional variability across infant formulas. (A) Overview of the study cohort. (B) Acquisition and processing of formula composition data. (C) Coefficients of variation for formula components across 17 formula brands. Iron, highlighted in red, was selected for subsequent experimental validation. Components present in a single formula brand, including 3’-galactosyllactose, *L. fermentum* and *L. rhamnosus* GG, are not shown because no coefficients of variation could be calculated. (D) Experimental validation workflow for the *in vitro* assessment of bacterial growth responses to iron limitation.

Given the observed nutritional variability, we next examined the associations of individual formula components, with infant gut microbiome diversity and specific bacterial taxa. Among the components that were significantly associated with bacterial taxa (nominally or after FDR-correction), iron was selected for downstream experimental validation due to its considerable variability across formulas, the higher iron concentrations in formula compared to breast milk ^31^, and iron’s established role in resource competition within microbial communities ^32,33^ (Fig. 1D).

### Formula composition influences species richness in the infant gut microbiome

Using mixed linear regression models, we explored the associations between 51 binary-classified formula components and alpha diversity metrics over time. Association analysis between formula components and species richness revealed nine nominally significant associations, with two still significant after FDR correction (Fig. 2A): Iodine was positively correlated with species richness (Fig. 2B), while vitamin E displayed a negative correlation (Fig. 2C). Association with the Shannon diversity index yielded weak signals, with only two components (vitamin B6 and magnesium) showing nominally significant and negative effects (Fig. S2). Together these results suggest that formula components may influence the number of bacterial species supported in the infant gut, potentially by altering nutrient availability or resource limitation and thereby affecting opportunities for bacterial establishment and persistence.

**Fig. 2.**
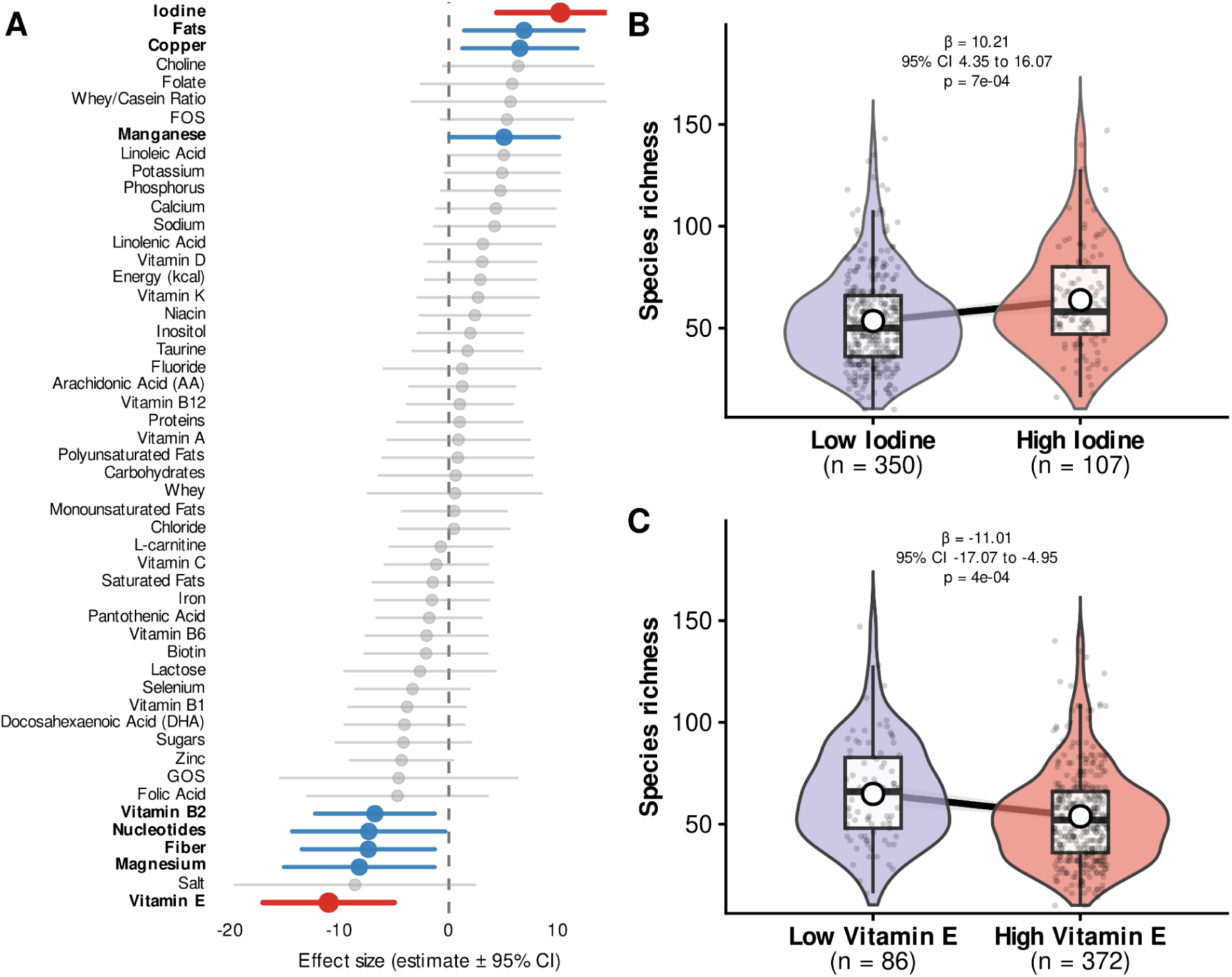
Formula composition is associated with species richness in the infant gut microbiome. (A) Effect sizes of binary-classified formula components on microbial species richness. Points show mixed-effects model estimates. Horizontal lines indicate 95% confidence intervals. Components with an FDR-adjusted *p* < 0.05 are shown in red, nominally significant associations (*p* < 0.05) are shown in blue, and non-significant associations shown in grey. (B,C) Species richness in infants consuming formulas with low or high concentrations of iodine (B) and vitamin E (C), pooled across the first three months of life. Violin plots show the distribution of observed richness. Boxplots show the median and interquartile range. Points represent individual samples. Black lines show the model-estimated change in richness from the low- to high-concentration group, with grey shading indicating the 95% confidence interval of the fitted mean and white circles indicating fitted values. Model coefficients (β) for species richness as a predictor, 95% confidence intervals, and *p*-values are shown above each panel.

### Formula components select for distinct bacterial taxa

We observed robust associations between individual formula components and specific bacterial species. Using mixed linear regression models, we tested the relationship between binary-classified formula components and the variation of CLR-transformed species’ abundances over time. Across 3,570 tests, 355 associations reached nominal significance and 26 remained significant after FDR correction. Here we found inverse associations between certain components and the abundance of specific *Bifidobacterium* species and other potentially opportunistic taxa. Specifically, *B. breve* and *B. bifidum* were negatively associated with the relative amount of vitamin D, zinc, iron, biotin, selenium and energy (kcal) and positively associated with phosphorus, fructo-oligosaccharides (FOS), sodium and calcium (Fig. 3). Interestingly, certain species within the class Clostridia (*Clostridium paraputrificum*, *Clostridium perfringens*, and *Intestinibacter bartlettii*) displayed inverse correlation patterns, showing increased abundance in response to components negatively associated with Bifidobacteria, including iron, selenium, and biotin, while being negatively correlated with the FOS abundance in the infant formula (Fig. 3).

**Figure 3.**
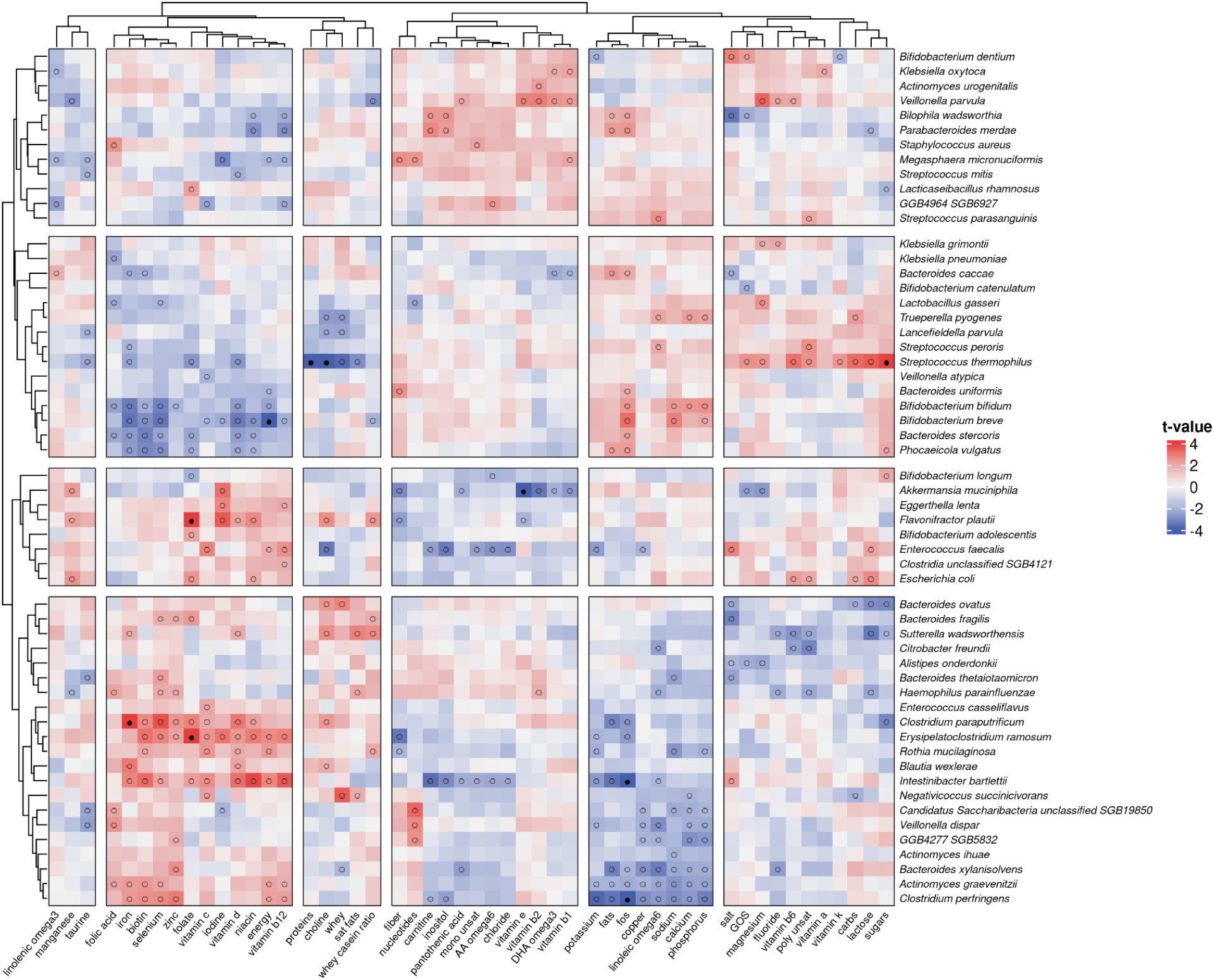
Formula components are associated with distinct species-level patterns in the infant gut microbiome. Hierarchically clustered associations between binary-classified infant formula components (columns) and bacterial species abundance (rows). Cell colour represents the *t*-value of the component–species association, with red indicating positive and blue indicating negative associations. Open circles indicate nominal significance (*p* < 0.05). Filled circles indicate associations significant after false-discovery-rate correction (FDR < 0.05). Only species and formula components with at least one nominally significant association are shown. Rows and columns are visually partitioned into four species clusters and six formula-component clusters, respectively.

Next, to model both the occurrence of species and validate results from the CLR-transformed association signals, we applied mixed logistic regression models to infer the presence or absence of species based upon binary-classified formula components. This analysis revealed 14 FDR-significant associations (Fig. S3), which confirmed previous signals. Iron concentration was negatively associated with *B. breve* presence (FDR = 0.003, Fig. S4A), while FOS concentration showed a positive association (FDR = 0.001, Fig. S3). Among the strongest negative predictors of species presence were zinc, iron, biotin, and selenium, which were significantly associated with the absence of *B. breve* and *B. bifidum* (Fig. S3). Conversely, linoleic acid, total fats, sodium, and FOS emerged as the strongest positive predictors, with FOS consistently associated with increased abundance and presence of *B. breve* and *B. bifidum* (Fig. 3, S3). Together, these results suggest that specific components can act as selective pressures that shape microbial colonization at the species level.

### Iron availability differentially shapes bacterial colonization

Iron emerged as a key modulator of microbial composition across our species abundance and presence-absence analyses. In the abundance models, lower iron concentrations were associated with increased abundances of *B. breve* and *B. bifidum* (Fig. 4). Logistic regression analysis confirmed that *B. breve* (FDR = 0.003, Figure S4B) and *B. bifidum* (log ratio transformation (LRT) p-value < 0.001, Figure S3) were more likely to be present in infants fed low iron formula. In contrast, higher formula iron levels were positively associated with the abundances of *C. paraputrificum* (estimate: 1.6, FDR = 0.03, Fig. 3 and Fig. S5A), *Clostridium perfringens* (estimate: 1.24, LRT p-value = 0.016, Fig. 3 and Fig. S5B), and *Intestinibacter bartlettii* (estimate: 1.2, LRT p-value = 0.005, Fig. 3 and Fig. S5C), suggesting that higher iron availability may promote the growth of these species. Taken together, these results suggest that iron availability exerts species-specific selective pressures on the infant gut microbiota, potentially favouring opportunistic taxa at the expense of beneficial *Bifidobacterium* species.

**Fig. 4.**
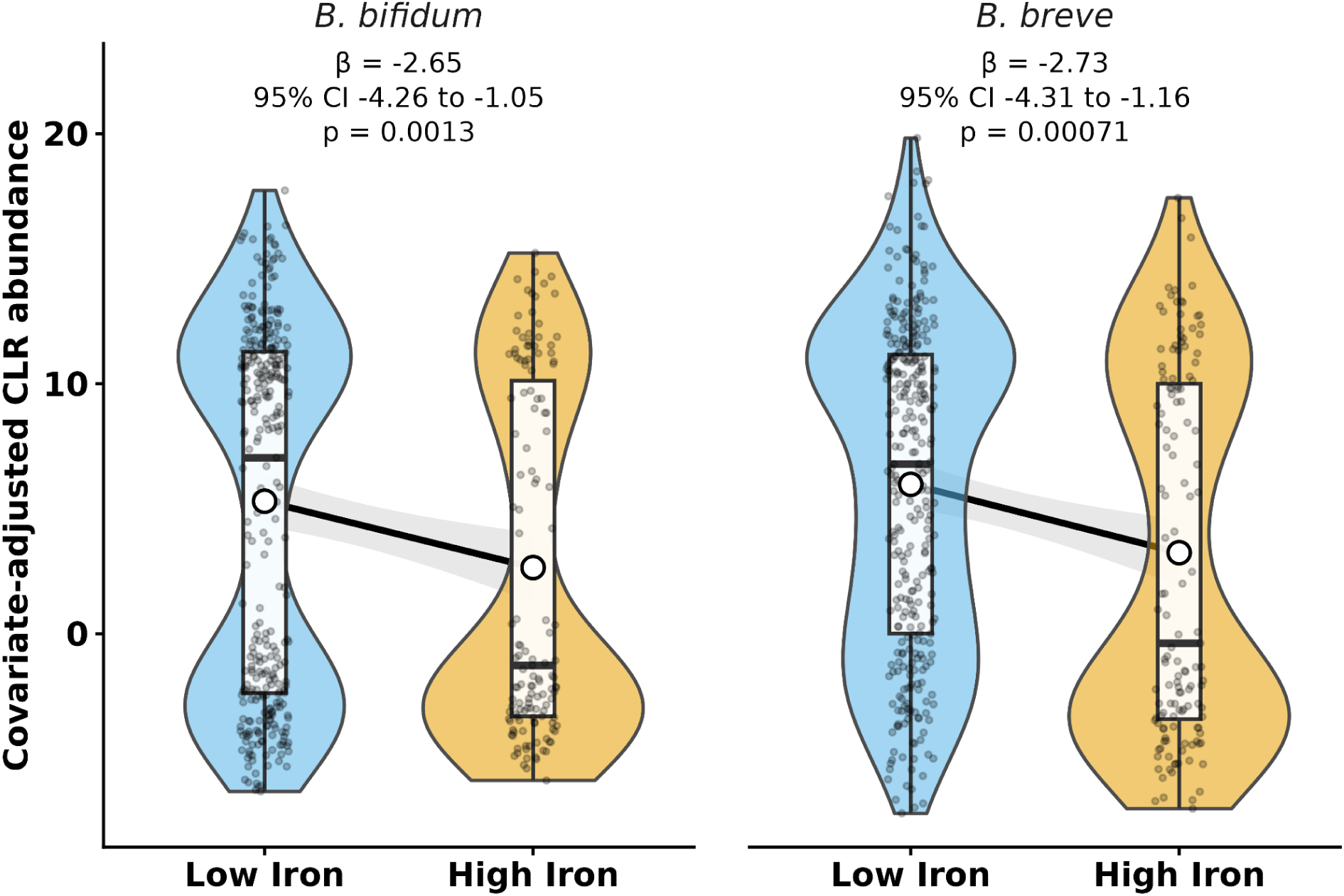
Iron concentration in infant formula is associated with *Bifidobacterium bifidum* and *Bifidobacterum breve* abundance. Boxplots show covariate-adjusted CLR-transformed relative abundance of *B. bifidum* and *B. breve* between low- and high-iron formula groups. CLR values were adjusted for time, birth mode, DNA concentration, sequencing batch, and sequencing depth using linear mixed-effects models, with infant ID included as a random intercept. Violin plots show the distribution of adjusted values, boxplots indicate the median and interquartile range, and points represent individual samples. Black lines indicate the model-estimated change from low- to high-iron formula. Shaded bands indicate 95% confidence intervals.

### Iron limitation reveals divergent phenotypic responses and their iron-related gene profiles

To complement the cohort associations, we performed exploratory growth assays to assess how selected infant-derived bacterial strains responded to iron chelation. Strains isolated from cohort stool samples were cultured with 0–70 μM of the iron chelator BPDS for 24 hours in triplicate. Under these conditions, *B. breve* and *B. bifidum* exhibited dose-dependent growth reduction, whereas *C. paraputrificum* showed complete growth inhibition at 30μM BPDS concentration (Fig. 5). These findings suggest that iron availability differentially affects bacterial taxa, and this may contribute to the associations we observed in the cohort.

**Fig. 5.**
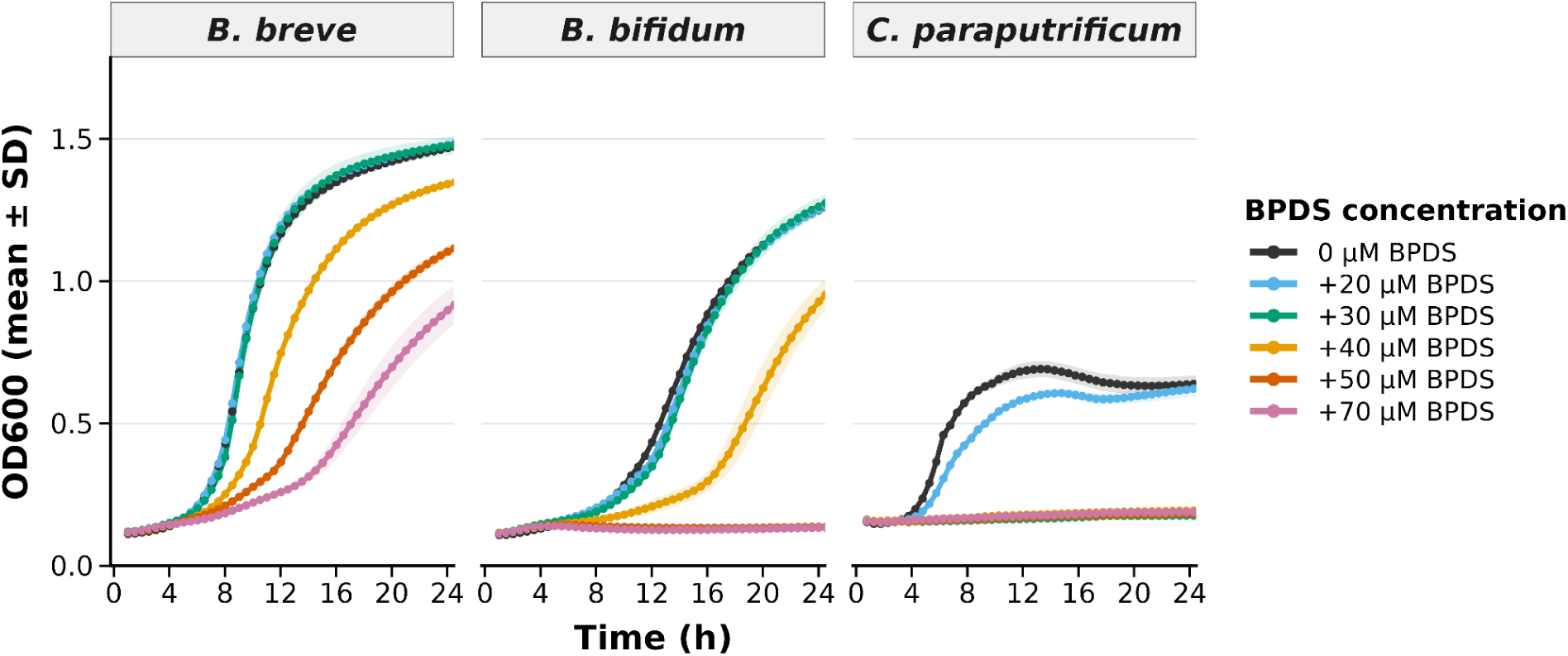
Iron chelation with BPDS alters bacterial growth *in vitro*. Strains of *Bifidobacterium breve*, *Bifidobacterium bifidum*, and *Clostridium paraputrificum* isolated from LLNEXT infant stool samples were cultured anaerobically overnight in YCFA lactose medium. Cultures were supplemented with BPDS at 0 (control), 20, 30, 40, 50 and 70 μM concentrations. Optical density measurements were taken every 30 min for a period of 24 hours, and growth curves were plotted for each strain. Data shown are mean optical density of triplicates ± standard deviation.

To explore the genetic basis underlying these phenotypic differences, we assessed iron-related genes in infant-associated *B. breve*, *B. bifidum* and *C. paraputrificum*. We observed distinct differences in iron gene profiles between the three species (Fig. S6), that potentially contribute to the divergent physiological responses detected under iron limitation. While a variety of iron-associated genes were detected across all three species, no specific iron uptake genes were detected in *C. paraputrificum* (Fig. S6). Notably, the well-characterized *bfeUOB* operon was detected in >94% of *B. bifidum* and *B. breve* genomes, and was completely absent in *C. paraputrificum* (Fig. S6). This operon, which encodes a high-affinity ferrous iron-uptake system that shares homology with the *efeUOB* transport system in *Escherichia coli*, has previously been shown to be highly upregulated in *B. breve* UCC2003 under iron-limiting conditions ^25^. The widespread conservation of *bfeUOB* across *Bifidobacterium* genomes suggests they have an enhanced iron-acquisition capacity that may contribute to *Bifidobacterium’s* resilience under iron limitation ^34^ and to the increased presence and abundance we observe in infants fed formulas with lower iron concentrations.

## Discussion

Research on the impact of formula feeding on the infant gut microbiome has largely focused on supplementation with selected bioactive ingredients, such as human milk oligosaccharides, leaving the impact of the broader variation in nutrient composition across formulas less explored. Here, we identified associations between the concentrations of specific formula components and microbial diversity and species composition in exclusively formula-fed infants. We focused on iron for exploratory follow-up based on its variability across formulas, its established role in bacterial growth and resource competition ^35,36^, and its contrasting associations with infant-associated Bifidobacteria and selected Clostridia.

Overall, we observed substantial differences in the concentrations of both macronutrients (e.g., proteins) and micronutrients including minerals (e.g., iron, iodine, fluoride) and vitamins (e.g., vitamin E) among the 17 infant formulas consumed by the LLNEXT infants. While carbohydrate and fat content was relatively consistent across formulas, over half of all formula components exhibited a CV > 25%, with fluoride exceeding 100%—a reflection of substantial variability across brands and the absence of a required minimum level under current EU regulations ^37^. These findings highlight the heterogeneity in nutritional exposure among formula-fed infants, which also appears to be reflected by the divergent alpha diversity and taxonomic associations observed across our analyses.

Key amongst our findings was that variation in formula components was better able to infer the level of species richness rather than Shannon diversity. This is because more components were either nominally or FDR-significant in associations to richness. In the case with richness, the effects of different components inferred both increased and decreased levels of richness. In contrast, only negative effects of two components were found when they were used to infer Shannon Diversity. Furthermore, the effect of components after multiple testing correction that were estimated via species richness, including vitamin E and Iodine, were indeed stronger than those effects of Shannon diversity. Iodine showed a positive association with richness, whereas vitamin E was negatively associated, suggesting that these components differentially shape ecological niche availability. Rather than acting as direct growth substrates, these compounds may function as environmental filters. Vitamin E, a potent antioxidant ^38^, may reduce oxidative heterogeneity within the gut environment, thereby favouring a narrower subset of taxa that are adapted to low oxidative stress conditions and limiting overall colonization. In contrast, higher iodine levels may reflect conditions in which fewer constraints limit microbial colonization, allowing a broader range of taxa to persist. Together, these findings support the idea that specific formula components influence gut colonization in formula-fed infants by modulating niche availability rather than altering dominance structure. Moreover, alleviation of competition for resources like iron, copper, and manganese, which all showed a nominally positive association with richness, may explain why formula-fed infants have previously exhibited a higher alpha diversity than breastfed infants ^11^.

Given that formula components appeared to modulate niche availability at community level, we examined species-level associations to determine how these environmental filters translate into species-specific responses. Here we observed numerous associations, indicating that component composition can selectively promote or suppress specific bacterial taxa in the infant gut. For instance, the prebiotic FOS was positively associated with *B. breve* and *B. bifidum,* consistent with previous studies showing that FOS supplementation promotes bifidobacterial growth in a dose- and duration-dependent manner ^35,36^. In contrast, higher iron concentrations were associated with the reduced abundance and presence of *B. breve* and *B. bifidum*, alongside high abundance of *C. paraputrificum*. Our exploratory growth assays provided additional context for these associations. Iron limitation reduced the growth of *Bifidobacterium breve and B. bifidum* but more strongly inhibited *C. paraputrificum*. Under iron-limited conditions, differences in iron requirements and acquisition strategies may influence the ability of bacteria to sustain growth and compete for available resources. Competition for iron therefore offers a plausible ecological framework for interpreting the contrasting cohort associations, although competitive interactions were not directly assessed. Other taxa positively associated with higher-iron formulas included *Clostridium perfringens,* which has been implicated in necrotising enterocolitis ^39,40^, and *Intestinibacter bartlettii,* which is typically rare in the infant gut but becomes more abundant as the microbiota matures ^41,42^. These findings align with studies showing that excess iron can disrupt microbial balance by promoting the growth of potentially pathogenic taxa, including *Clostridium spp.,* while suppressing beneficial Bifidobacteria ^43–45^.

This ecological interpretation also raises the question of how iron supply differs between feeding modes. Formula iron concentrations ranged from 3.3 to 12.1 mg/L in our cohort (mean = 6.6 mg/L), which stands in contrast to reported breast milk concentrations < 1 mg/L ^31^. Breast milk also contains iron-binding proteins such as lactoferrin ^46^, which highlights the importance of considering both iron concentration and its accessibility to bacteria. Our genomic analysis offers further context. The *bfeUOB* operon was detected in > 94% of the *B. breve* and *B. bifidum* genomes analyzed, but it was not detected in *C. paraputrificum*. In contrast, *C. paraputrificum* possesses a substantially larger repertoire of iron-associated genes (median = 29) than *B. bifidum* (median = 7) or *B. breve* (median = 8), indicating distinct iron-acquisition strategies. All in all, greater iron availability may favour taxa such as *C. paraputrificum* and *C. perfringens* by easing iron competition. Alongside iron, the other formula components associated with microbial diversity and species composition represent candidate nutritional factors for further investigation to inform formula optimization.

In summary, our study shows that infant formula composition varies widely and that formula components may influence the trajectory of the early-life gut microbiota. While overall microbial alpha diversity was moderately affected by the variability in formula components, species-level associations revealed underlying competition dynamics through which individual components could shape microbial populations. However, several limitations should be acknowledged. First, detailed data on feeding frequency and quantity per feeding were not available. These factors vary between infants and depend on individual weight and caregiver interpretation of feeding cues, thereby introducing variability into our analysis that could influence microbial outcomes. Second, we could not fully account for changes in formula type over-time. Although only 12% of infants switched formulas between timepoints, this could still introduce variability in nutrient exposure and microbial response.

More broadly, this study offers insights into the relationship between infant formula composition and early-life gut microbiota development. We show that micronutrients such as iron and vitamin E could affect the colonization patterns of the infant microbiome. Given the variation in infant formula composition and the naturally occurring variation in breast milk composition, and the differences between them, future research should examine how this variability shapes early microbial colonization, immune development, and long-term health.

## Resource availability

### Lead contact

Requests for further information and resources should be directed to and will be fulfilled by the lead contact, Cyrus A. Mallon.

### Materials availability

This study did not generate new, unique reagents.

### Data and code availability

All original code has been deposited on Github and is publicly available. The bioinformatic pipeline for microbiome data processing is available at: (https://github.com/GRONINGEN-MICROBIOME-CENTRE/gmc-mgs-pipeline)

The bioinformatic pipeline for data preparation and association analyses is available at: (https://github.com/GRONINGEN-MICROBIOME-CENTRE/Lifelines_NEXT/tree/main/Formula_Project). Detailed instructions for reproducing the analyses are provided in the repository’s README file.

### Consortia

Lifelines NEXT cohort study authors and affiliations

Milla F. Brandão-Gois^1^, Marcel Bruinenberg^2^, Siobhan Brushett^1,3^, Jackie A.M. Dekens^1,4^, Sanzhima Garmaeva^1^, Sanne J. Gordijn^5^, Soesma A. Jankipersadsing^1^, Ank de Jonge^6,7^, Gerard H. Koppelman^8,9^, Marlou L.A. de Kroon^3^, Folkert Kuipers^8,10^, Alexander M. Kurilshikov^1^, Anouk Marsman^2^, Lilian L. Peters^6,7^, Jelmer R. Prins^5^, Sijmen A. Reijneveld^3^, Sicco Scherjon^5^, Jan Sikkema^4^, Trishla Sinha^1^, Johanne E. Spreckels^1^, Aline B. Sprikkelman^8,9^, Morris A. Swertz^1^, Henkjan J. Verkade^8^, Cisca Wijmenga^1^, Alexandra Zhernakova^1^

^1^ Department of Genetics, University of Groningen and University Medical Center Groningen, Groningen, the Netherlands

^2^ Lifelines Cohort Study and Biobank, Groningen, the Netherlands

^3^ Department of Health Sciences, University of Groningen and University Medical Center Groningen, Groningen, the Netherlands

^4^ Innovation Center, University Medical Center Groningen, Groningen, the Netherlands

^5^ Department of Obstetrics and Gynecology, University of Groningen and University Medical Center Groningen, Groningen, the Netherlands

^6^ Department of Primary and Long-term Care, University of Groningen, University Medical Center Groningen, Groningen, the Netherlands

^7^ Midwifery Science, Amsterdam University Medical Center, Vrije Universiteit Amsterdam, AVAG, Amsterdam Public Health, Amsterdam, The Netherlands

^8^ Department of Pediatrics, University of Groningen and University Medical Center Groningen, Groningen, the Netherlands

^9^ Groningen Research Institute for Asthma and COPD (GRIAC), University of Groningen and University Medical Center Groningen, Groningen, the Netherlands

^10^ European Research Institute for the Biology of Ageing (ERIBA), University of Groningen and University Medical Center Groningen, Groningen, the Netherlands

## Acknowledgments

The data used in this manuscript are provided by LLNEXT. The authors are grateful for the participation of all the parents and infants in LLNEXT. We thank the LLNEXT team and maternity care providers for recruiting participants, collecting material during and after childbirth, and building and maintaining the cohort. We also thank K. Mc Intyre for English and content editing and the Genomics Coordination Center and the Center for Information Technology of the University of Groningen for their support and for providing access to the Gearshift, Peregrine, and Habrok high-performance computing clusters. The Lifelines NEXT cohort study received funds from the University Medical Center Groningen Hereditary Metabolic Diseases Fund, Health∼Holland (Top Sector Life Sciences and Health), the EU, the Northern Netherlands Alliance (SNN), the provinces of Friesland and Groningen, the municipality of Groningen, Philips, and the Société des Produits Nestlé. This work was supported by funds from the EASI-Genomics grant (PID7780) to T.S. and A.Z. T.S. holds a scholarship from the Junior Scientific Masterclass, University of Groningen and a De Cocks-Hadders Stichting grant (Winston Bakker Fonds WB-08). A.Z. is supported by the Nederlandse Organisatie voor Wetenschappelijk Onderzoek (NWO) VIDI grant VI.C.232.074, the EU Horizon Europe Program grant INITIALISE (101094099), NWO KIC grant KICH1.LWV04.21.01, and NWO Gravitation grant Exposome-NL (024.004.017).

## Author contributions

A.Z. and C.A.M. conceived the project. A.K. performed the bioinformatic analysis for the microbiome data processing. T.S. processed microbiome data and feeding and delivery information in LLNEXT. J.A.D.D. performed the analysis of iron-related genes in infant-associated metagenome-assembled genomes. I.F. performed the association analysis, under the supervision of C.A.M. I.F.,Y.J., and T.K. performed the lab experiments, under the supervision of C.A.M. I.F., C.A.M., and A.Z. were involved in data interpretation. I.F. prepared the first draft of the manuscript and led the writing process, all other authors contributed to the editing of the manuscript.

## Declaration of Generative AI Use

Generative AI was used to assist with coding for data analysis and manuscript writing. The authors reviewed and verified all AI-assisted code. The authors take full responsibility for the accuracy and integrity of the analyses and the manuscript.

