## Supplementary Information for "Infant formula composition is linked to distinct patterns of early gut colonization"

### Supplementary Figures

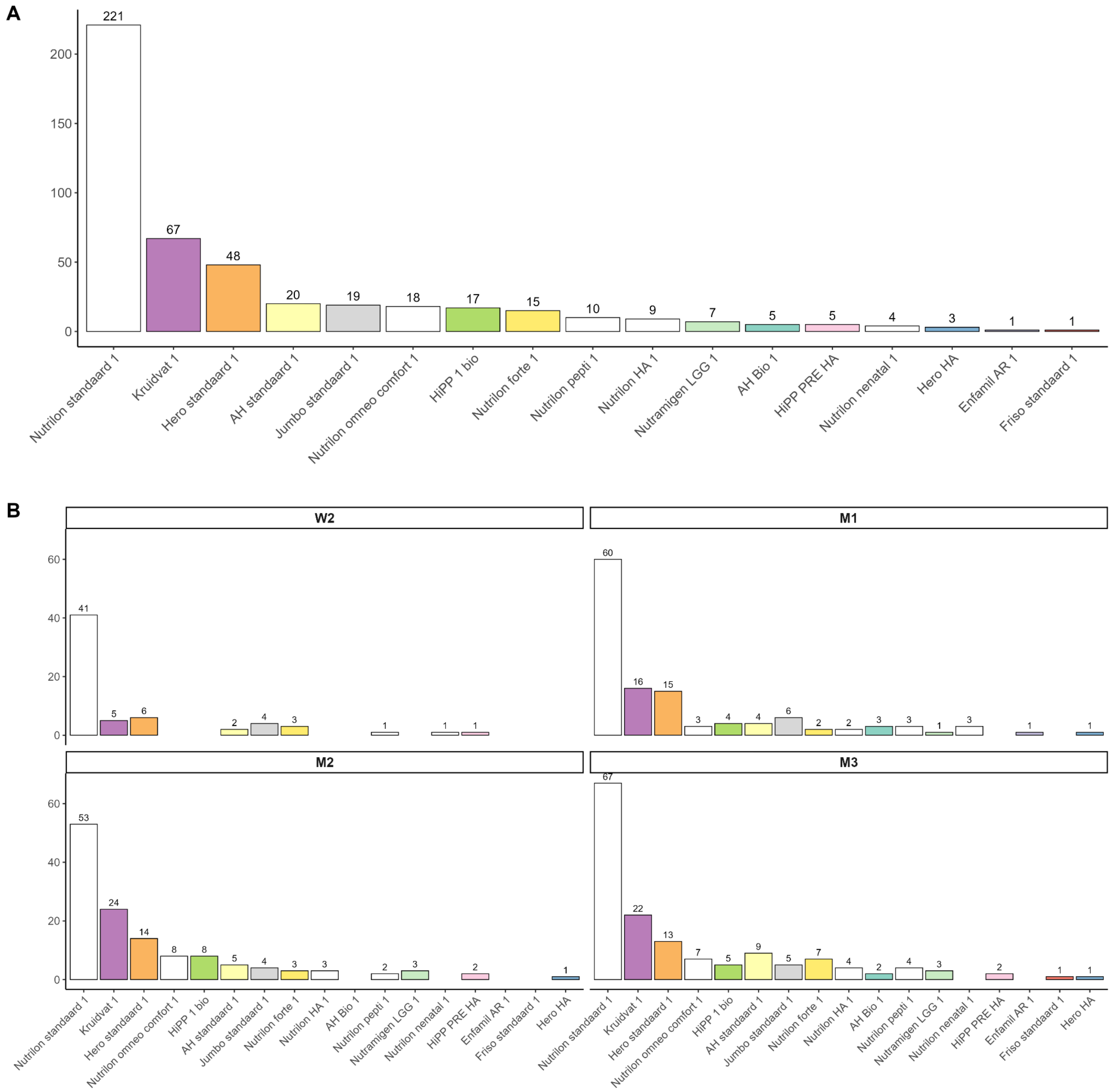


**Figure S1. Sample distribution across formula brands and timepoints.** (A) Total number of samples collected for each formula across all timepoints (W2–M3). (B) Number of samples assigned to each formula at individual timepoints. Each panel represents a different timepoint and shows the frequency of each formula given.

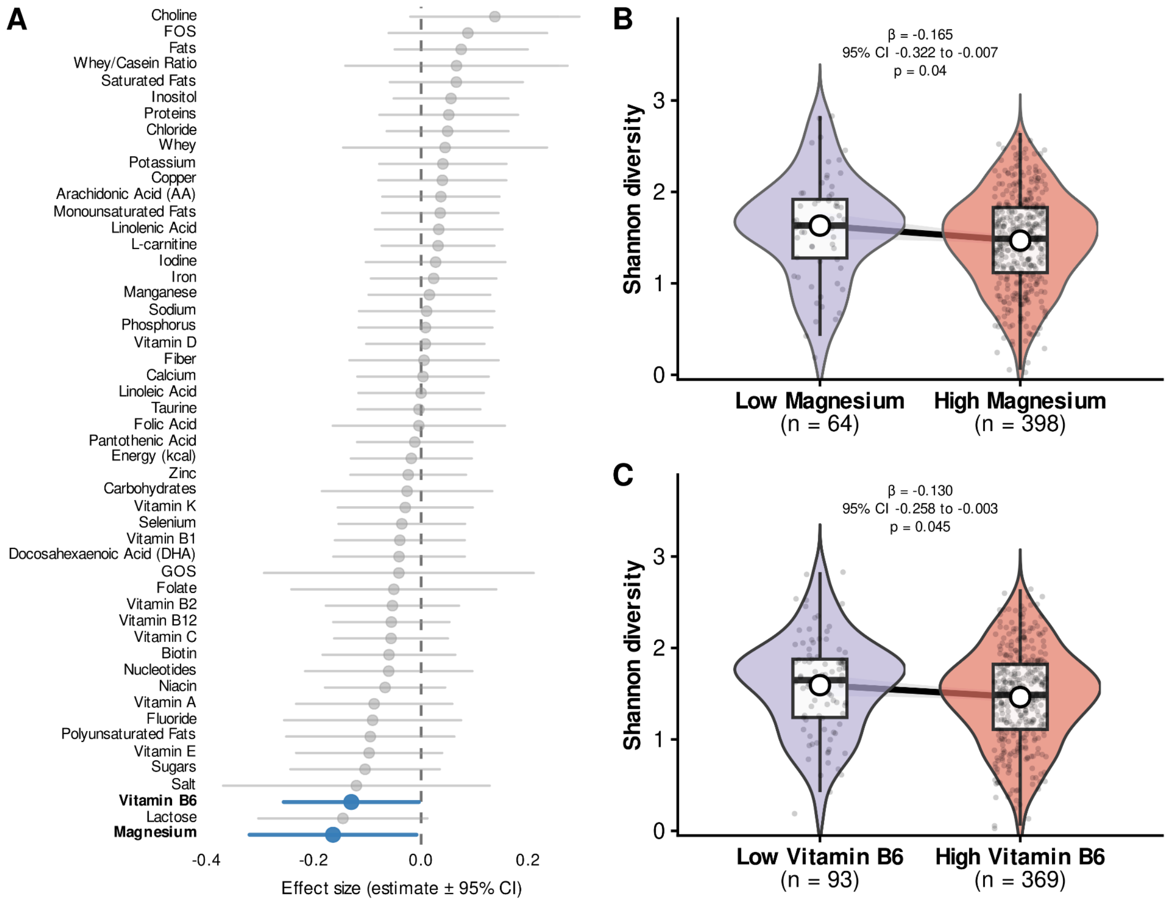


**Figure S2. Shannon diversity is not strongly associated with formula composition.** (A) Effect sizes of binary-classified formula components on Shannon diversity index, with 95% confidence intervals. Significance is indicated as nominal (likelihood-ratio test of the mixed linear regression model, cutoff 0.05). (B,C) Boxplots and violin-plots comparing microbiome Shannon diversity index between low- and high-concentration groups for magnesium (B) and vitamin B6 (C), pooled across all timepoints. Significance is based on the likelihood-ratio test without FDR correction. * p-value <0.05. The individual sample points are also shown.

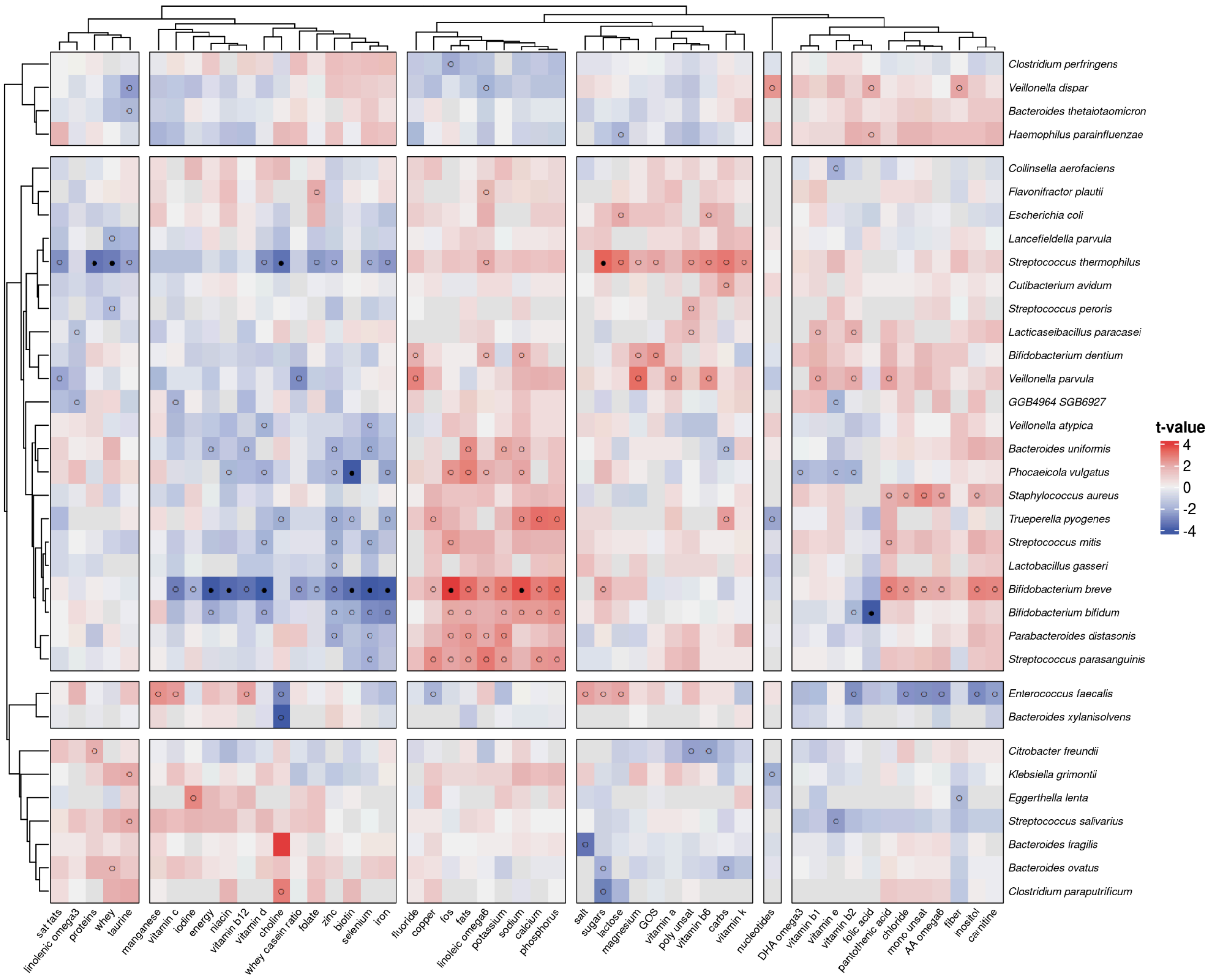


**Figure S3. Heatmap for associations between the component groups with bacterial species presence.** Results are derived from a mixed logistic regression model. Colours represent the t-value of the associations. Open circles indicate nominal significance ($p < 0.05$). Filled circles indicate associations significant after false-discovery-rate correction (FDR < 0.05)

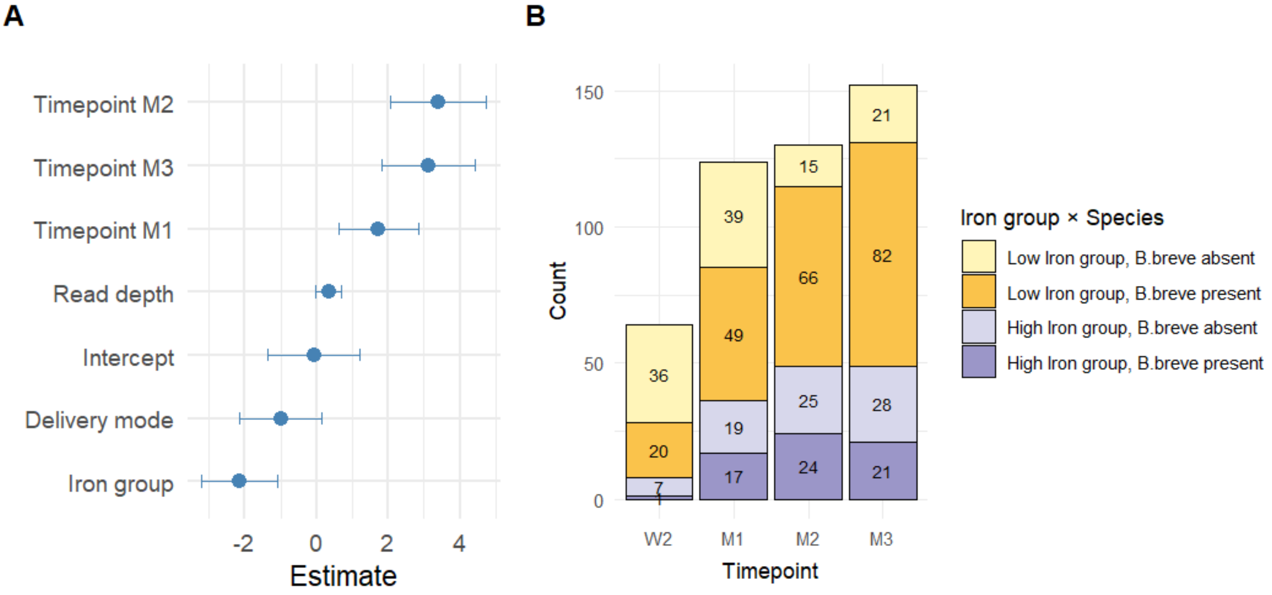


**Figure S4. FDR-significant association between iron formula group and *Bifidobacterium breve* presence.** (A) Logistic regression estimates for predictors of *B. breve* presence. Error bars represent 95% confidence intervals. (B) Distribution of samples with *B. breve* present or absent across low- and high-iron formula groups.

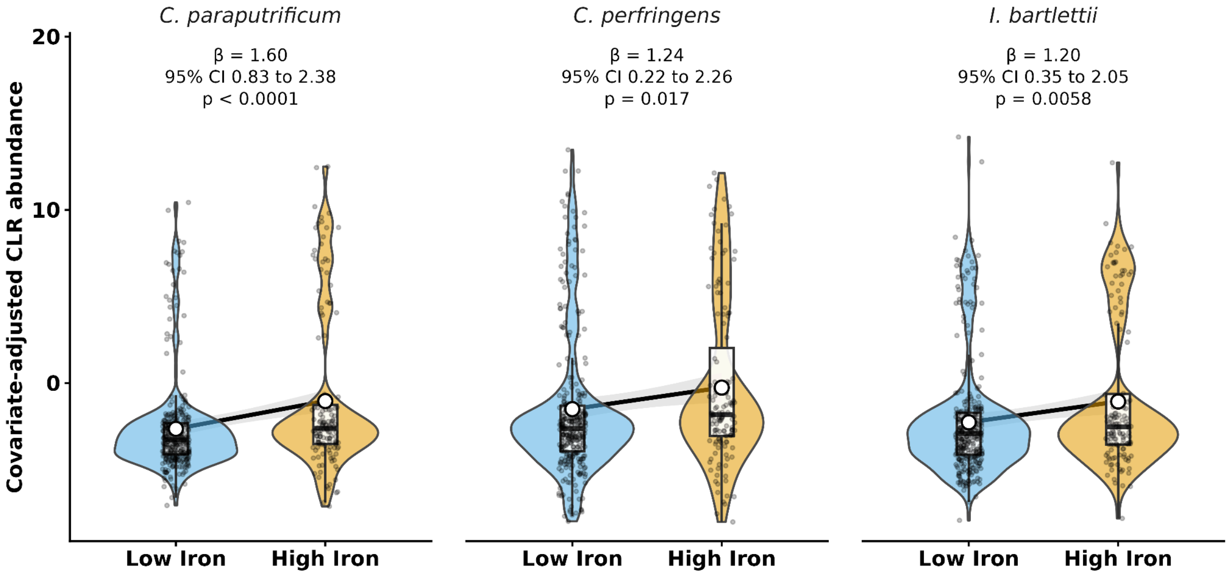


**Figure S5. Boxplots showing CLR-transformed relative abundance of *Clostridium paraputrificum* (A), *Clostridium perfringens* (B), and *Intestinibacter bartlettii* (C) across all timepoints, grouped by iron formula group.** Iron groups were defined based on whether the formula's iron concentration was below/equal to the median value (low iron) or above (high iron). For *C. paraputrificum,* the p-value is the FDR-adjusted result from the likelihood ratio test of the mixed linear regression model. For *C. perfringens* and *I. bartlettii*, the p-values are unadjusted likelihood-ratio test results of mixed linear regression models (nominal significance).

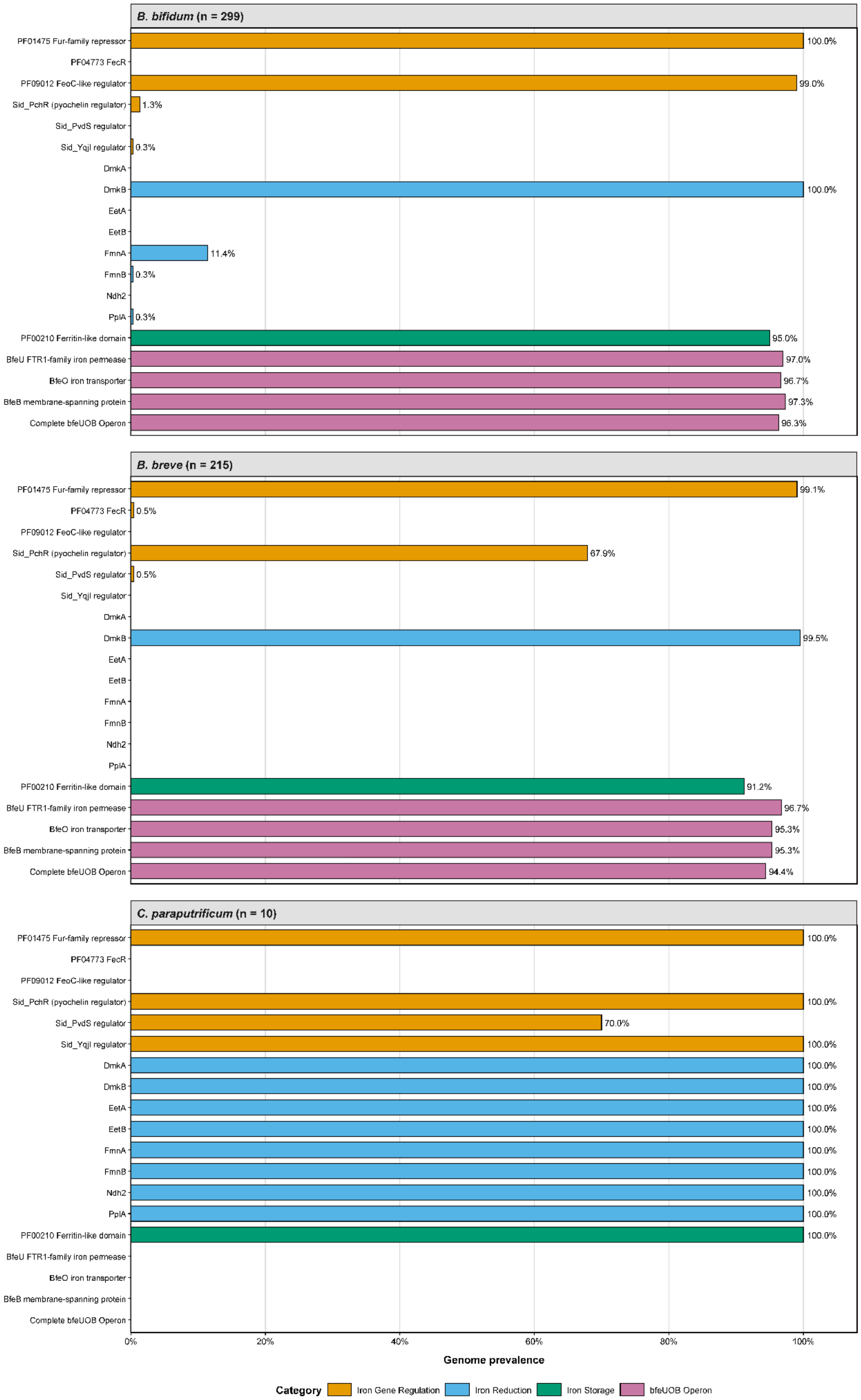

B. bifidum (n = 299)
B. breve (n = 215)
C. paraputrificum (n = 10)
PF01475 Fur-family repressor
PF04773 FecR
PF09012 FeoC-like regulator
Sid_PchR (pyochelin regulator)
Sid_PvdS regulator
Sid_YqjI regulator
DmkA
DmkB
EetA
EetB
FmnA
FmnB
Ndh2
PplA
PF00210 Ferritin-like domain
BfeU FTR1-family iron permease
BfeO iron transporter
BfeB membrane-spanning protein
Complete bfeUOB Operon
100.0%
99.0%
1.3%
0.3%
100.0%
11.4%
0.3%
0.3%
95.0%
97.0%
96.7%
97.3%
96.3%
99.1%
0.5%
67.9%
0.5%
99.5%
91.2%
96.7%
95.3%
95.3%
94.4%
100.0%
100.0%
70.0%
100.0%
100.0%
100.0%
100.0%
100.0%
100.0%
100.0%
100.0%
100.0%
100.0%
0%
20%
40%
60%
80%
100%
Genome prevalence
Category
Iron Gene Regulation
Iron Reduction
Iron Storage
bfeUOB Operon

**Figure S6. Summary of the prevalence of iron-related genes detected in infant-associated *B. breve*, *B. bifidum*, and *C. paraputrificum.*** Horizontal bars represent the prevalence (%) of iron-related genes across metagenome-assembled genomes of each species. Colours indicate the predicted functional categories: iron gene regulation (orange), iron reduction (blue), iron storage (green), and the iron uptake operon *bfeUOB* (pink).
